# Role of 5-HT2A, 5-HT2C, 5-HT1A and TAAR1 receptors in the head twitch response induced by 5-hydroxytryptophan and psilocybin: Translational implications

**DOI:** 10.1101/2022.07.22.501026

**Authors:** Orr Shahar, Alexander Botvinnik, Noam Esh-Zuntz, Michal Brownstien, Rachel Wolf, Gilly Wolf, Bernard Lerer, Tzuri Lifschytz

## Abstract

There is increasing interest in the therapeutic potential of psilocybin in psychiatric disorders. In common with other serotonergic psychedelics, psilocybin is thought to act via the 5-HT2A receptor (5-HT2AR). Serotonin is the endogenous ligand of 5-HTR. In rodents, the serotonin precursor, 5-hydroxytryptophan (5-HTP), and psilocybin, induce a characteristic head twitch response (HTR), which is correlated with the human psychedelic trip in intensity and duration. We examined the role of other serotonergic receptors and the trace amine associated receptor 1 (TAAR1) in modulating HTR induced by 5-HTP and psilocybin. Male C57BL/6J mice (11 weeks old, ~30g) were administered 5-HTP, 50-250 mg/kg intraperitoneally (i.p.) or 200 mg/kg i.p. after pretreatment with 5-HT/TAAR1 receptor modulators. Psilocybin was administered at 0.1-51.2 mg/kg i.p. or at 4.4 mg/kg i.p. preceded by 5-HT/TAAR1 receptor modulators. HTR was assessed in a custom-built magnetometer. 5-HTP and psilocybin induced a dose dependent increase in the frequency of HTR over 20 minutes with attenuation by the 5-HT2AR antagonist, M100907 (volanserin), and the 5-HT1AR agonist, 8-OH-DPAT. The 5-HT2CR antagonist, RS102221, enhanced HTR at lower doses but reduced it at higher doses for 5-HTP and psilocybin. The TAAR1 antagonist, EPPTB, reduced 5-HTP-but not psilocybin-induced HTR. We have confirmed the key role of 5-HT2AR in HTR and have demonstrated an effect of 5-HT1AR and a bimodal contribution of 5-HT2CR as well as a role of TAAR1 in modulating HTR induced by 5-HTP. Compounds that modulate HTR induced by psychedelics have a potentially important role in the emerging therapeutic use of these compounds.

**Significance Statement:** We have confirmed the key role of 5-HT2AR in in the induction of HTR by 5-HTP and psilocybin, have demonstrated the effect of a 5-HT1AR agonist to attenuate HTR and a bimodal contribution of 5-HT2CR as well as a role of TAAR1 in modulating HTR induced by 5-HTP. Compounds that modulate HTR induced by psychedelics have a potentially important role in the emerging therapeutic use of these compounds.

**Visual Abstract:** 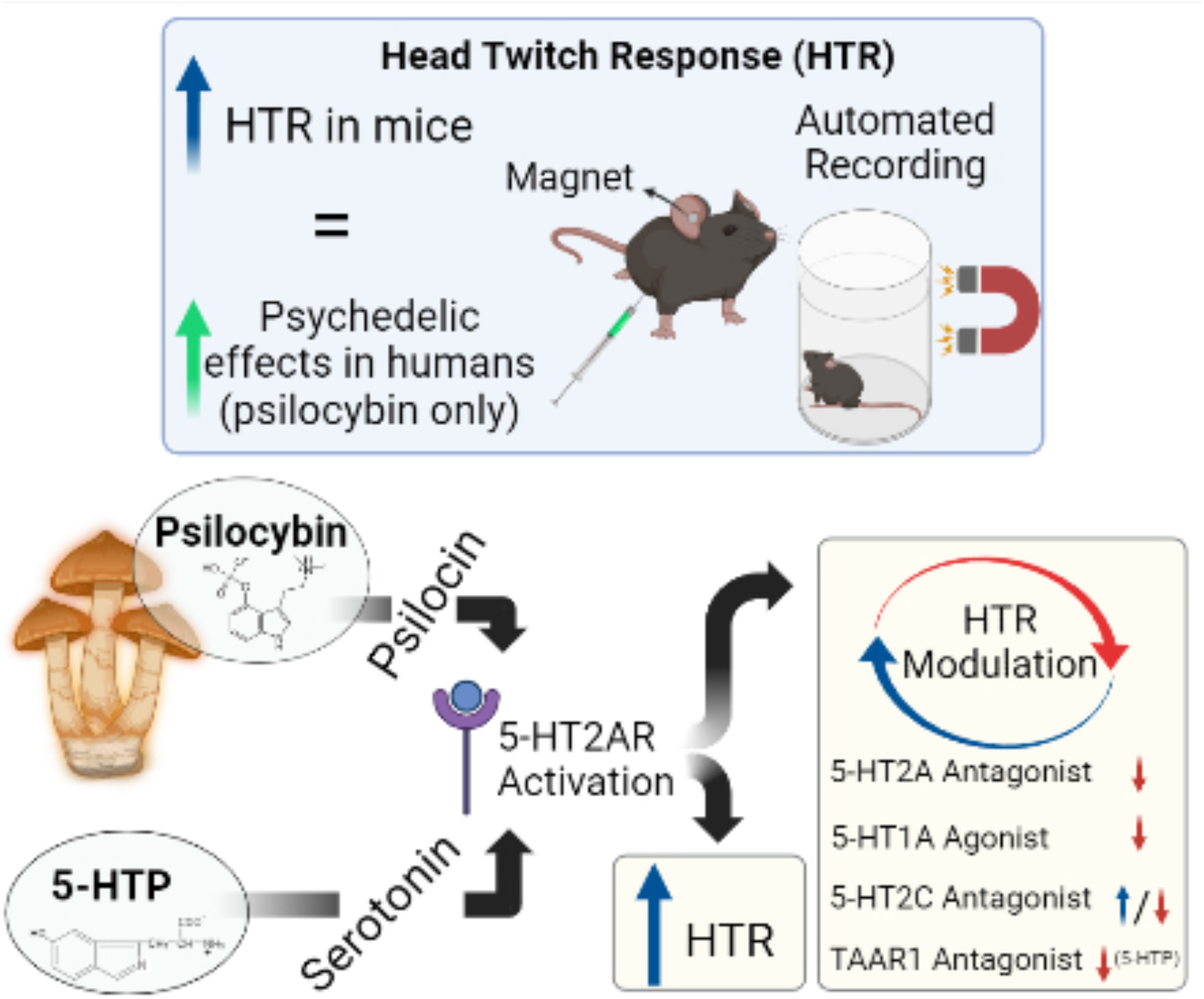

## Introduction

Psychiatric disorders cause significant individual suffering as well as having a major economic impact (Knapp and Wong, 2020; Castelpietra et al., 2022) and manifest a high level of resistance to standard treatment (Gershkovich et al., 2017; Cepeda et al., 2018). Novel therapeutic approaches are needed. Recent findings suggest that psychedelic compounds may fill this role. Psychedelics significantly alter consciousness, cognition and mood (Carhart-Harris and Goodwin, 2017; Nutt, 2022) and have been used for centuries for healing and spiritual purposes (Garcia-Romeu and Richards, 2018; Winkelman, 2019). After he synthesized lysergic acid diethylamide (LSD) in 1938 and discovered its psychedelic properties, Albert Hoffman synthesized psilocybin which is the principal psychoactive component of “magic mushrooms” (Hofmann et al., 1959). A significant body of mostly uncontrolled research followed but was halted when psychedelic drugs were declared Schedule 1 substances in 1968 (Nutt, 2022). After decades of relative inactivity, research on psychedelic compounds has increased significantly, including controlled studies in depression (Carhart-Harris et al., 2021; Galvao-Coelho et al., 2021) and posttraumatic stress disorder (Krediet et al., 2020; Mitchell et al., 2021). A number of trials are ongoing (Siegel et al., 2021). The principal emphasis has been on serotonergic psychedelics which include tryptamines such as psilocybin, N,N-dimethyltryptamine (DMT) and 5-methoxy-N,N-dimethyltryptamine (5-MeO-DMT); phenethylamines such as mescaline; and ergolines such as LSD.

Serotonergic psychedelic agents such as psilocybin are thought to act mainly via the serotonin 5-HT2A receptor to which psilocybin’s active metabolite, psilocin, binds with high affinity (Nichols, 2016; Kyzar et al., 2017). Although it is known that serotonergic psychedelics bind to an array of serotonin receptors including but not limited to 5-HT1A/2A/2B/2C (Slocum et al., 2022), it is not fully understood how activation of different serotonin receptors interacts to exert the subjective and neurobiological effects of psychedelics (Passie et al., 2002).

In rodents, psilocybin induces a characteristic head twitch response (HTR), which is correlated with the psychedelic trip in humans and across rodent species (Halberstadt et al., 2020). Because of increasing interest in the development of novel psychedelic analogues, including variants that do not induce the characteristic “trip” (Cameron et al., 2021; Lu et al., 2021), HTR is an important preclinical tool in the evaluation of psychedelic compounds. HTR was first described in mice after administration of the serotonin precursor, 5-hydroxytryptophan (5-HTP) (Corne et al., 1963) and has been further characterised by subsequent investigators (González-Maeso et al., 2003; Fantegrossi et al., 2005; Fantegrossi et al., 2006; González-Maeso et al., 2007; Keiser et al., 2009; Fantegrossi et al., 2010; Halberstadt, 2020). Although extensive research has documented the effect of 5-HTP to induce HTR in rodents (Darmani and Reeves, 1996; Sun et al., 2003; Devi and Sharma, 2014; Zhuk et al., 2015), psychedelic effects have not been reported at doses administered to humans (de la Fuente Revenga et al., 2021).

Studies in humans have shown that 5-HT2A antagonists, such as ketanserin can block the short-term subjective effects of psilocybin and other serotonergic psychedelics (Vollenweider et al., 1998; Olbrich et al., 2021). Attenuation of subjective psychedelic effects has also been reported for the 5-HT1A partial agonist, buspirone (Pokorny et al., 2016). It is well established that 5-HT2A antagonists block HTR in mice administered psilocybin and other serotonergic psychedelics (Hesselgrave et al., 2021). Fantegrossi et al. (2010) have shown that the 5-HT2C antagonist, RS102221, enhances DOI-elicited-HTR in C57Bl/6 mice at doses up to 3 mg/kg and attenuates HTR at a dose of 10 mg/kg. Custodio et al. (2022) showed substantial attenuation of HTR induced by DOI and two derivatives of mescaline by pre-treatment with the 5-HT2C antagonist, SB-242084. A very interesting finding was that one of the mescaline derivatives induced HTR that was blocked by the 5-HT2C antagonist, SB242084, but not by a 5-HT2A antagonist (ketanserin), indicating that the HTR-inducing effect was mediated by 5-HT2CR alone (Custodio et al., 2022).

In the current study, we sought to define the dose-response characteristics of HTR induced by 5-HTP and chemically synthesized psilocybin (PSIL) in C57Bl/6j mice, to examine the role of 5-HT2A, 5-HT2C and 5-HT1A receptors in modulating HTR induced by 5-HTP and PSIL using appropriate agonists and antagonists and to determine a possible role of the trace amine associated receptor 1 (TAAR1) in mediating HTR induced by 5-HTP and psilocybin. Our findings confirm the pivotal role of 5-HT2A receptors in HTR induced by 5-HTP and PSIL, support an inhibitory role of 5-HT1A receptors, and an inverted U doseresponse effect of a 5-HT2C receptor antagonist and suggest a role for TAAR1 receptors in the HTR-inducing effect of 5-HTP.

## Methods

### Animals

Experiments were performed on adult (9–12 weeks old) C57BL/6J male mice. Animals were housed under standardized conditions with a 12-h light/dark cycle, stable temperature (22 ± 1 °C), controlled humidity (55 ± 10 %) and free access to food and water. Experiments were conducted in accordance with AAALAC guidelines and were approved by the Authority for Biological and Biomedical Models, Hebrew University of Jerusalem, Israel, Animal Care and Use Committee. All efforts were made to minimize animal suffering and the number of animals used.

### Drugs

PSIL was supplied by Usona Institute, Madison, WI, USA and was determined by AUC at 269.00 nm (UPLC) to contain 98.75% psilocybin. 5-HTP, M100907, 8-OH-DPAT, and EPPTB were purchased from Sigma-Aldrich, Israel. RS102221 was purchased from Biotest, Israel. PSIL was dissolved in 100% saline (0.9% NaCl) solution. 5-HTP was dissolved in 5% DMSO + 95% saline (0.9% NaCl) solution. M100907, 8-OH-DPAT, and RS102221 were dissolved in a 5% DMSO + 95% saline (0.9% NaCl) solution. EPPTB was dissolved in 5% ethanol + 5% kolliphor + 90% saline (0.9% NaCl) solution. All solutions were prepared to the appropriate volume of 10 μl/g and concentration for administration (by intraperitoneal [i.p.] injection). Vehicle-treated condition represents injection of the appropriate solvent to the equivalent volume of the drug administered.

### Ear tagging

Assessment by electromagnetic generation of the rapid side-to-side headshake that characterizes HTR requires the installation of small magnets in the outer ears of the mice. For this purpose, we utilized small neodymium magnets (N50, 3 mm diameter × 1 mm height, 50 mg) which were attached to the top surface of aluminum ear tags (supplied by Mario de la Fuente Revenga PhD. of Virginia Commonwealth University). Ear tags were placed through the pinna antihelix and laid in the interior of the antihelix, resting on top of the antitragus, leaving the ear canal unobstructed. This procedure was performed by simple restraint and immobilization of the mouse’s head. Signs of ear tissue damage were rare in the form of redness, and the ear tags were well tolerated in the mice.

### Head Twitch Acquisition

Mice were allowed to recover from ear-tagging for 5-7 days prior to testing. The tagged animals were placed inside a magnetometer apparatus (supplied by Mario de la Fuente Revenga PhD. of Virginia Commonwealth University, Supplementary Fig. 1A) consisting of plastic containers (11.6 cm diameter X 13.3 cm height) surrounded by a coil (~500 turns 30 AWG enameled wire) the output of which was amplified (Pyle PP444 phono amplifier) and recorded at 1000 Hz using a NI USB-6001 (National Instruments, US) data acquisition system (de la Fuente Revenga et al., 2020). Recordings were performed using a MATLAB driver (MathWorks, US, R2021a version, along with the NI myDAQ support package) with the corresponding National Instruments support package for further processing. A custom MATLAB script was used to record the processed signal, which was presented as graphs showing the change in current as peaks (mAh). A custom graphic user interface created in our laboratory was used to further process the recording into an Excel spreadsheet.

For validation of the magnetometer recording a total of 5 trials that included treatments of vehicle (n=1), PSIL 0.75 (n=1), PSIL 2.2 mg/kg (n=2), and PSIL 4.4 mg/kg (n=1) were administered with parallel magnetometer and video recording (Supplementary Fig. 1C, Supplementary Fig. 2). A Hero Black 9 GoPro camera (GoPro) was used to record high frame rate (120 frames per sec) overhead videos (1080p resolution). GoPro camera was mounted ~ 18 cm above the magnetometer coil floor, and experimental test sessions were recorded for 30 min post injection. After videos of each experiment were recorded, the video files were transferred to hard drive for storage and subsequent visual analysis. Visual scoring verified that HTRs obtained by magnetometer recording were the same peaks seen visually by the observer (Supplementary Fig. 1B). The validation shows a perfect linear regression line (R^2^ = 1) of the visual scoring and magnetometer recording. Out of 307 magnetometer peaks that were validated, only 1 was found to be a false positive on visual examination. There were no HTRs that a human observer found that the magnetometer did not record.

Drugs were administered by intraperitoneal (i.p.) injection immediately before placing the weighed animals into the magnetometer. For dose response, 5-HTP and PSIL were administered at doses of 50 mg/kg to 250 mg/kg and 0.1 mg/kg to 51.2 mg/kg, respectively. For assessing the effects of modulators, all mice were injected with either 5-HTP or PSIL at doses of 200 and 4.4 mg/kg respectively preceded by the 5-HT2A receptor antagonist M107900 (0.5, and 2 mg/kg i.p.), the 5-HT2C receptor antagonist RS102221 (2, 4, and 8 mg/kg i.p.), the 5-HT1A receptor agonist 8OH-DPAT (1 and 2 mg/kg i.p.) or the TAAR 1 antagonist EPPTB (1 and 10 mg/kg i.p.). HTR was measured for 30 minutes for 5-HTP and 20 minutes for PSIL in the magnetometer.

### Statistical analysis

The experimental data are expressed as the mean ± standard deviation (SD). To determine inter group differences, one- and two-way repeated measure analysis of variance (ANOVA) were used, with or without repeated measures, as indicated. Tukey’s and Dunnett’s Multiple Comparison Test was used to analyze post-hoc comparisons. p < 0.05 (two tailed) was the criterion for significance. Graph Pad Prism, version 9.3.1 software was used for all statistical analyses.

## Results

### Effect of 5-HTP on HTR

Initially, we evaluated the effect of increasing doses (50 mg/kg to 250 mg/kg) of 5-HTP on HTR. These 5-HTP doses induced a dose dependent increase in HTR (Fig. 1A) reflected in a between-subject main effect of dose in a two-way repeated measure ANOVA (F [5, 23] = 8.328, p = 0.0001). HTR rate increased during the test, indicated by within-subject effect of time (F [3.830, 77.09] = 9.051, p < 0.0001), and by time x dose interaction (F [70, 322] = 2.031, p < 0.0001). When comparing total HTR (Fig. 1B), there was a notable increase of HTR at the higher doses (One-way ANOVA: F [5,23] = 8.328, p = 0.0001; post hoc: Vehicle vs. 5-HTP 150 mg/kg p = 0.0343, Vehicle vs. 5-HTP 200 mg/kg p = 0.0002, Vehicle vs. 5-HTP 250 mg/kg p = 0.0204). 5-HTP induced the greatest HTR (Fig. 1A) during the last 15 min of the 30 min measurement. We therefore decided it would be most relevant to analyse the effects of receptor modulators on 5-HTP induced HTR during the 15-30 min measurement period for the following experiments.

**Figure 1.**
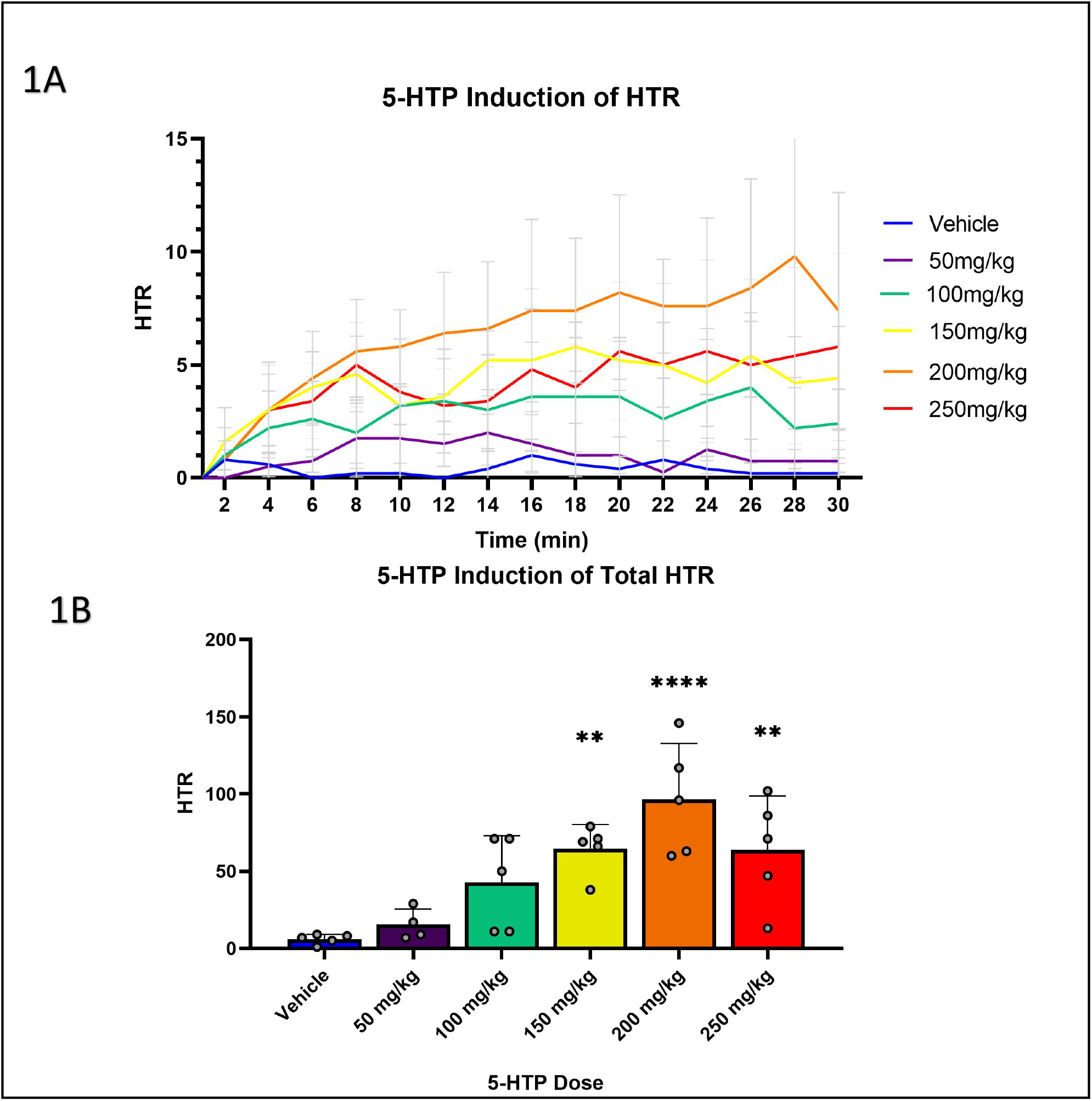
Induction of HTR by 5-HTP, over time (A), and total HTR during 30 min (B). (A) Effect of different 5-HTP concentrations during HTR over the time course of 30 min post-injection (n=4-5). Two-way ANOVA (A): Time F [3.830, 77.09] = 9.051, P < 0.0001. Dose F [5, 23] = 8.328, P = 0.0001. Time x Dose F [70, 322] = 2.031, P < 0.0001. (B) Cumulative HTR during the time of 30 min post injection (n=4-5). One-way ANOVA: F [5, 23) = 8.328, P = 0.0001. Dunnett post hoc. Compared to Vehicle, ** *P* < 0.01, *** *P* < 0.0001. Error bars represent SD.

To examine the effects of serotonin and TAAR1 receptor modulators, 5-HTP 200 mg/kg was administered following a dose of the modulators (Fig. 2). The 5-HT2A receptor agonist, M100907 (Fig. 2A) completely blocked 5-HTP induced HTR at both 0.5 mg/kg and 2 mg/kg doses (One-way ANOVA: F [2,14] = 15.65, p = 0.0002; Dunnett’s post hoc: Vehicle vs. M100907 0.5 mg/kg p = 0.0007, Vehicle vs. M100907 2 mg/kg p = 0.0004). Pre-treatment with the 5-HT1A receptor agonist, 8-OH-DPAT 1 mg/kg and 2 mg/kg (Fig. 2B) significantly reduced 5-HTP-induced HTR (One-way ANOVA: F [2,36] = 8.385, p = 0.0010; Dunnett’s post hoc: Vehicle vs. 8-OH-DPAT 1 mg/kg p = 0.0035, Vehicle vs. 8-OH-DPAT 2 mg/kg p = 0.0014). The 5-HT2C antagonist, RS102221 (Fig. 2C), did not alter 5-HTP induced HTR at 2 mg/kg, and significantly increased 5-HTP induced HTR at 4 mg/kg and 8 mg/kg (One-way ANOVA: F [3,13] = 7.229, p = 0.0042; Tukey’s post hoc: Vehicle vs. RS102221 4 mg/kg p = 0.0059, Vehicle vs. RS102221 8 mg/kg p = 0.0176). The TAAR1 antagonist, EPPTB (Fig. 2D) significantly reduced 5-HTP induced HTR at both doses of 1 mg/kg, and 10 mg/kg (One-way ANOVA: F [2,11] = 4.939, p = 0.0295; Dunnett’s post hoc: Vehicle vs. EPPTB 1 mg/kg p = 0.0421, Vehicle vs. EPPTB 10 mg/kg p = 0.0344).

**Figure 2.**
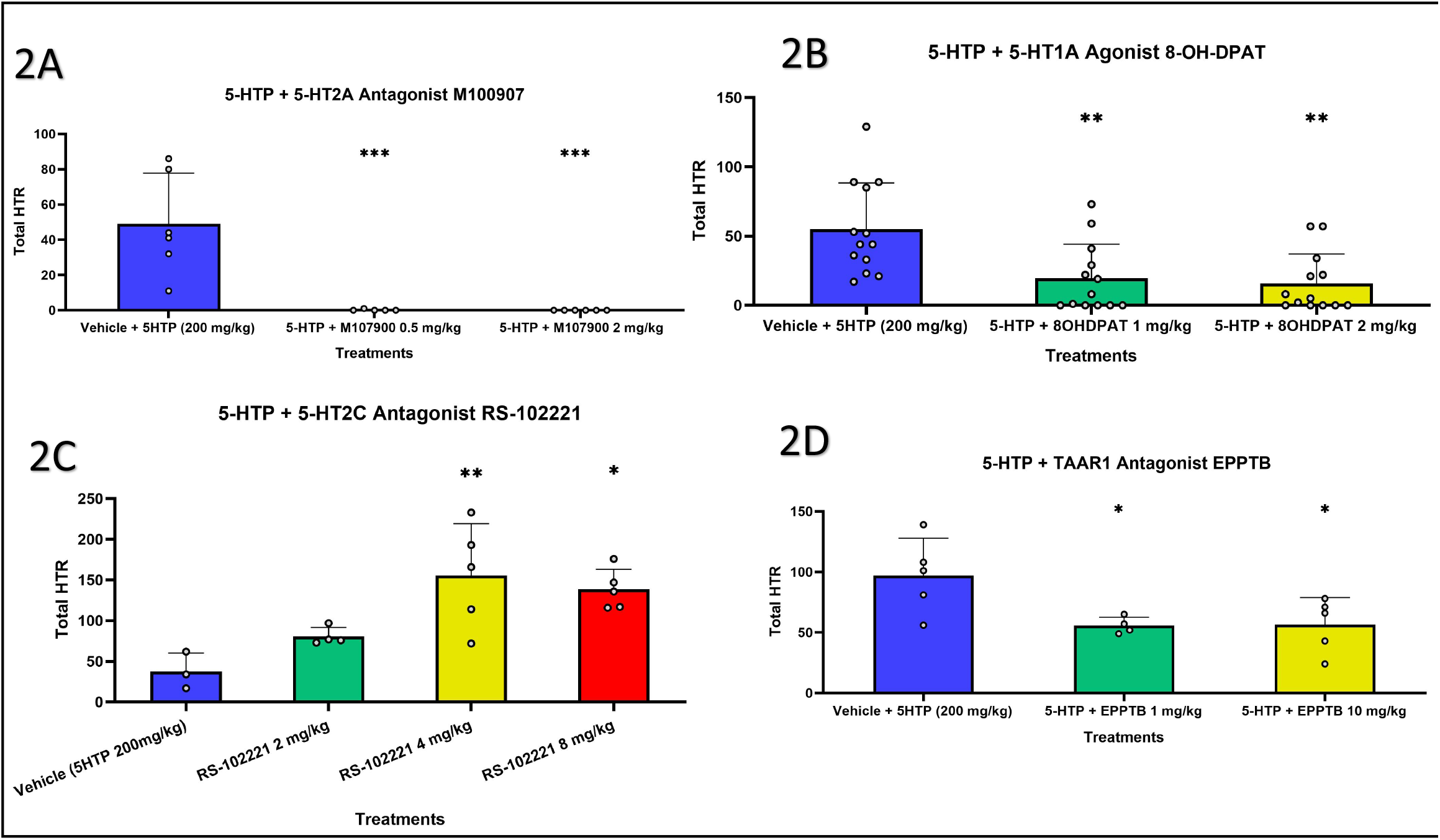
Total 5-HTP induced HTR (A-D) during 15-30 min post injection with different co-treatments. (A) Effect of pre-treatment with M100907 0.5 mg/kg +5-HTP 200 mg/kg, M100907 2 mg/kg + 5-HTP 200 mg/kg, or Vehicle + 5-HTP 200 mg/kg (n=5-6). (B) Effect of pre-treatment with 8-OH-DPAT 1 mg/kg + 5-HTP 200 mg/kg, 8-OH-DPAT 2mg/kg + 5-HTP 200 mg/kg, or Vehicle + 5-HTP 200mg/kg (n=13). (C) Effect of pre-treatment with RS102221 2 mg/kg + 5-HTP 200 mg/kg, RS102221 4 mg/kg + 5-HTP 200 mg/kg, RS102221 8 mg/kg + 5-HTP 200 mg/kg, or Vehicle + 5-HTP 200 mg/kg (n=3-5). (D) Effect of pre-treatment with EPPTB 1 mg/kg + 5-HTP 200 mg/kg, EPPTB 10mg/kg + 5-HTP 200 mg/kg, or Vehicle + 5-HTP 200 mg/kg (n=4-5). One-way ANOVA: (A) F [2, 14] = 15.65, P = 0.0003 Dunnett’s post hoc test. (B) F [2, 36] = 8.385, P = 0.0010 Dunnett’s post hoc test. (C) F [3, 13] = 7.229, P = 0.00.42 Tukey’s post hoc test. (D) F [2, 11] = 4.939, P = 0.0295 Dunnett’s post hoc test. Compared to Vehicle, * *P* < 0.05, ** *P* < 0.01, *** *P* < 0.001.

### Effect of Psilocybin on HTR

First, we assessed the effect of increasing PSIL concentrations (0.1 mg/kg to 51.2 mg/kg) on mouse HTR. HTR over time (2 min time bins) showed a dose dependent increase of head twitch up to a dose of 25.6 mg/kg and a reduced effect with the next increment dose of 51.2 mg/kg. (Fig. 3A) reflected in a between-subject main effect of dose in a two-way repeated measure ANOVA (F [16, 88] = 7.194, p < 0.0001). HTR rate increased during the test, indicated by within-subject effect of time F [3.309, 291.2] = 46.56, p < 0.0001), and by time x dose interaction (F [144, 792] = 8.838, p < 0.0001). The dose response graph was observed to manifest a bimodal profile (Fig. 3B). The lower PSIL doses from 0-1.6 mg/kg produced a gradual increase followed by a slightly sustained HTR and then gradual decrease of the effect when compared to the higher doses from 3 mg/kg – 51.2 mg/kg, that produced a very sharp onset with a fast decline of HTR without a sustained period. The doses were compiled into two groups (0.1 – 1.6 mg/kg, and 3 – 51.2 mg/kg) and analysed (Two-way ANOVA: Time F [2.2821, 262.4] = 58.25, P < 0.0001. Dose F [1, 93] = 15.39, P = 0.0002. Time x Dose F [9, 837] = 85.70, P < 0.0001). Total PSIL induced HTR during the 20 min measuring period was compiled (Fig. 3C); the doses that induced the most HTR over time when compared to vehicle were 1.6 mg/kg, 3.2 mg/kg, 4.4 mg/kg, and 6.5 mg/kg (One-way ANOVA: F [16,88] = 7.156, p < 0.0001; post hoc: Vehicle vs. PSIL 0.5 mg/kg p = 0.0398, Vehicle vs. PSIL 1.6 mg/kg p < 0.0001, Vehicle vs. PSIL 3 mg/kg p = 0.0032, Vehicle vs PSIL 3.2 mg/kg p < 0.0001, Vehicle vs PSIL 4.4 mg/kg p < 0.0001, Vehicle vs PSIL 6.4 mg/kg p < 0.0001, Vehicle vs PSIL 12.8 mg/kg p = 0.0374, Vehicle vs PSIL 25.6 mg/kg p = 0.0002).

**Figure 3.**
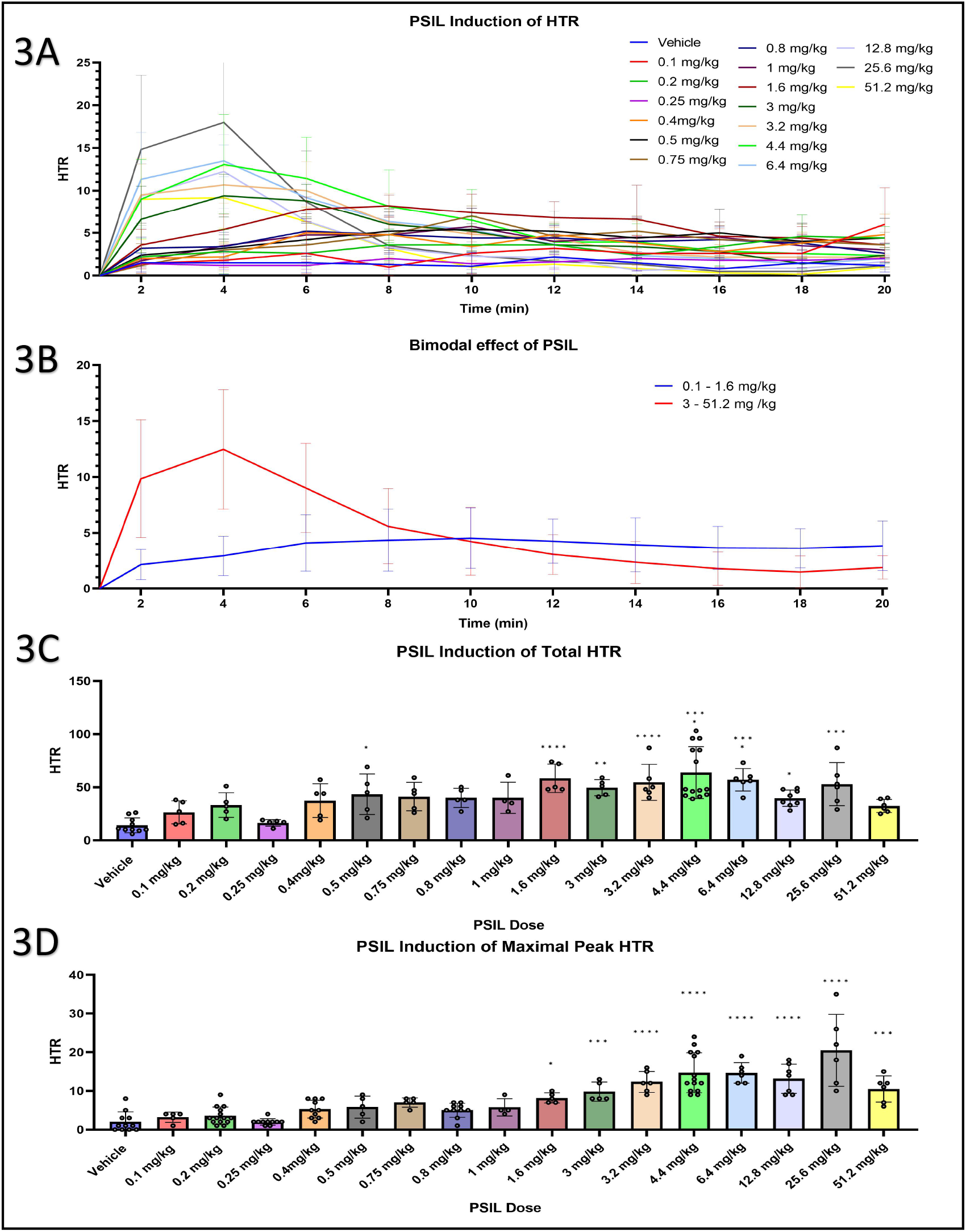
PSI induced HTR, over time (A), bimodal effect (B), total HTR during 20 min (C) and maximal peak HTR in a 2 min bin over 20 min (D) (n=5-14). (A) Effect of different PSIL concentrations during HTR over the time course of 20 min post-injection. Twoway ANOVA: Time F [3.309, 291.2] = 46.56, P < 0.0001. Dose F [16, 88] = 7.194, P < 0.0001. Time x Dose F [144, 792] = 8.838, P < 0.0001. (B) Bimodal HTR effect of high and low doses over the time course of 20 min post-injection. Two-way ANOVA: Time F [2.2821, 262.4] = 58.25, P < 0.0001. Dose F [1, 93] = 15.39, P = 0.0002. Time x Dose F [9, 837] = 85.70, P < 0.0001. (C) Cumulative HTR during the 20 min post injection. One-way ANOVA: F [16, 88] = 7.156, P < 0.0001. Tukey’s post hoc test. (D) Maximal peak HTR that was produced in a 2 min time bin over the course of 20 min. One-way ANOVA: F [16, 112] = 19.02, P < 0.0001. Tukey’s post hoc test. Compared to Vehicle, * *P* < 0.05, ** *P* < 0.01, *** *P* < 0.001, **** *P* < 0.0001. Error bars represent SD.

An important aspect of the evaluation was to examine the maximal effect that PSIL had on inducing HTR (Fig. 3D), which was determined through peak HTR comparison. When compared to vehicle, a significant difference was shown for the higher doses (above 1.6 mg/kg) in a similar fashion to the bimodal pattern seen in Fig. 3A (One-way ANOVA: F [16,112] = 19.02, p < 0.0001; post hoc: Vehicle vs. PSIL 3 mg/kg p = 0.0051, Vehicle vs. PSIL 3.2/4.4/6.5/12.8/25.6 mg/kg p < 0.0001, Vehicle vs. PSIL 51.2 mg/kg p = 0.0004). 25 mg/ 70 kg is administered as a representative high psilocybin dose for human clinical trials (Brown et al., 2017; van Amsterdam and van den Brink, 2022). Using DoseCal: a virtual calculator for dosage conversion between human and different animal species (Janhavi et al., 2022), 25 mg/70 kg human dose was converted to 4.4 mg/kg in mice. Administration of 4.4 mg/kg PSIL stimulated a strong acute HTR response (Fig. 3A). Although this dose was administered i.p. and is thus not analogous to an oral dose in humans, it chosen as a representative acute PSIL dose for further receptor modulator experiments.

To examine the effects of serotonin and TAAR1 receptor modulators, PSIL 4.4 mg/kg was administered following doses of the modulators (Fig. 4): 20 min post injection of PSIL, M100907 (Fig. 4A) completely blocked HTR at both 0.5 mg/kg and 2 mg/kg doses (One-way ANOVA: F [2,21] = 54.20, p < 0.0001; Dunnett’s post hoc: Vehicle vs. M100907 0.5 mg/kg p < 0.0001, Vehicle vs. M100907 2 mg/kg p < 0.0001). 20 min post injection of PSIL, treatment with 8-OH-DPAT (Fig. 4B) significantly reduced PSIL induced HTR at 1 mg/kg and considerably more so with 2 mg/kg (One-way ANOVA: F [2,20] = 19.53, p < 0.0001; Dunnett’s post hoc: Vehicle vs. 8-OH-DPAT 1 mg/kg p = 0.0003, Vehicle vs. 8-OH-DPAT 2 mg/kg p < 0.0001). 20 min post injection of PSIL, treatment with RS102221 (Fig. 4C) did not alter PSIL induced HTR with 2 mg/kg, significantly increased PSI induced HTR with 4 mg/kg and reduced PSIL induced HTR with 8 mg/kg (One-way ANOVA: F [3,24] = 8.178, p = 0.0006; Tukey’s post hoc: Vehicle vs. RS102221 4 mg/kg p = 0.0363; RS102221 2 mg/kg vs. RS102221 4 mg/kg p = 0.0371; RS102221 4 mg/kg vs. RS102221 8 mg/kg p = 0.0006). EPPTB (Fig. 4D) did not alter (no statistical sig.) PSIL induced HTR with both doses of 1 mg/kg and 10 mg/kg.

**Figure 4.**
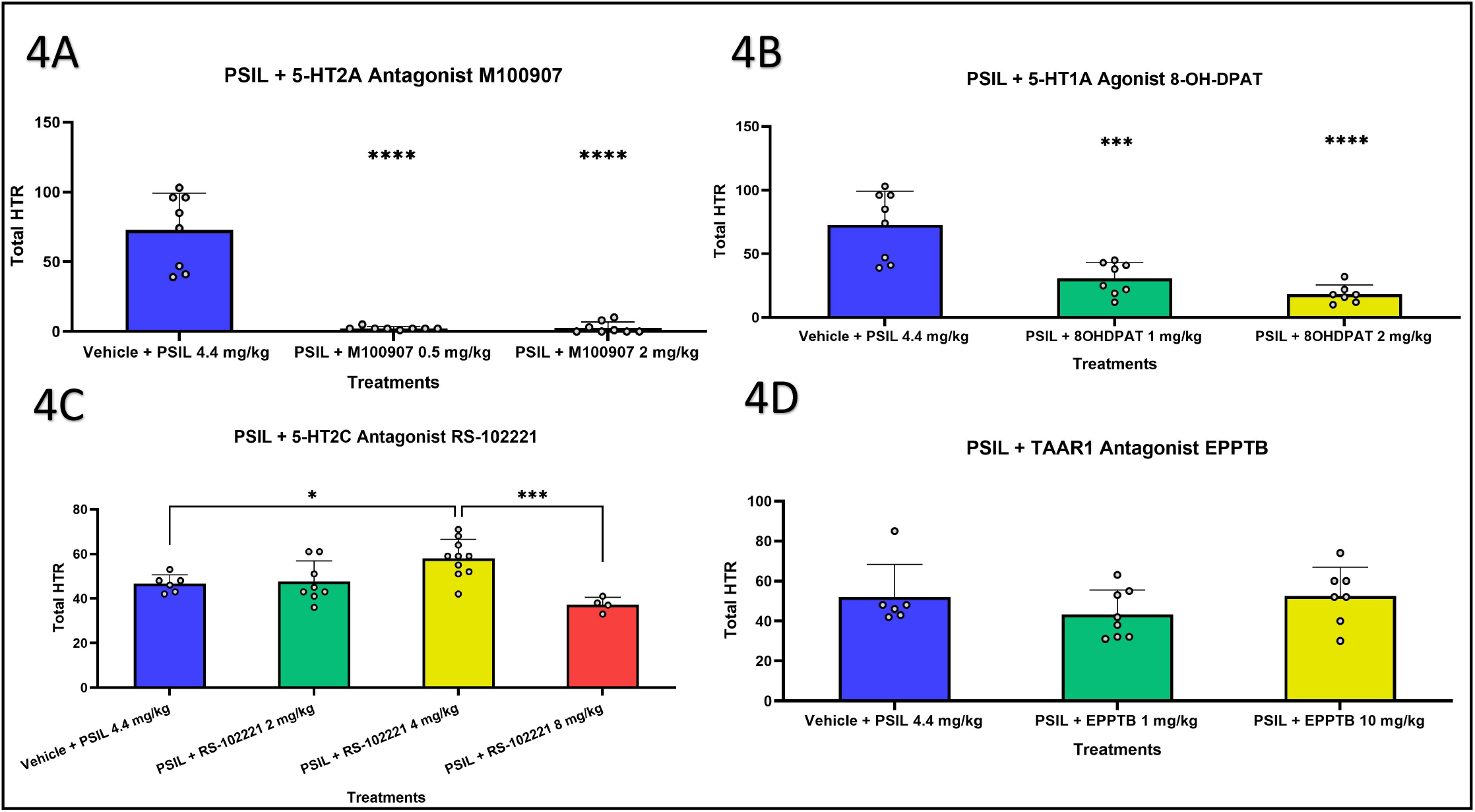
Total PSIL induced HTR (A-D) during 20 min post injection with different co-treatments. (A) Effect of pre-treatment with M100907 0.5 mg/kg + PSIL 4.4 mg/kg, M100907 2mg/kg + PSIL 4.4 mg/kg, or Vehicle + PSIL 4.4 mg/kg (n=8) (B) Effect of pre-treatment with 8-OH-DPAT 1 mg/kg + PSIL 4.4 mg/kg, 8-OH-DPAT 2mg/kg + PSIL 4.4 mg/kg, or Vehicle + PSIL 4.4 mg/kg (n=7-8) (C) Effect of pre-treatment with RS102221 2 mg/kg + PSIL 4.4 mg/kg, RS102221 4 mg/kg + PSIL 4.4 mg/kg, RS102221 8 mg/kg + PSIL 4.4 mg/kg, or Vehicle + PSIL 4.4 mg/kg (n=4-11) (D) Effect of pre-treatment with EPPTB 1 mg/kg + PSIL 4.4 mg/kg, EPPTB 10mg/kg + PSIL 4.4 mg/kg, or Vehicle + PSIL 4.4 mg/kg (n=6-8) One-way ANOVA: (A) F [2, 21] = 54.20, P < 0.0001. Dunnett’s post hoc test. (B) F [2, 20] = 19.53, P < 0.0001. Dunnett’s post hoc test. (C) F [3, 24] = 8.178, P = 0.0006. Tukey’s post hoc test. (D) F [2, 18] = 1.010, P = 0.3841. Dunnett’s post hoc test. Compared to Vehicle (A, B, D), and between treatments (C), * *P* < 0.05, ** *P* < 0.01, *** *P* < 0.001, **** *P* < 0.0001.

To assess any difference between 5-HTP and PSIL induction profile of HTR, we chose doses with comparable maximal HTR. 5-HTP 200mg/kg was compared to PSIL 3 mg/kg (Fig. 5). A somewhat “mirror image” HTR profile was obtained. Although both doses of compounds reached the same peak HTR, the way the HTR was induced was different (Two-way ANOVA: Compound F [1, 8] = 6.099, p = 0.0387. Time x Compound F [14, 112] = 13.91, p < 0.0001).

**Figure 5.**
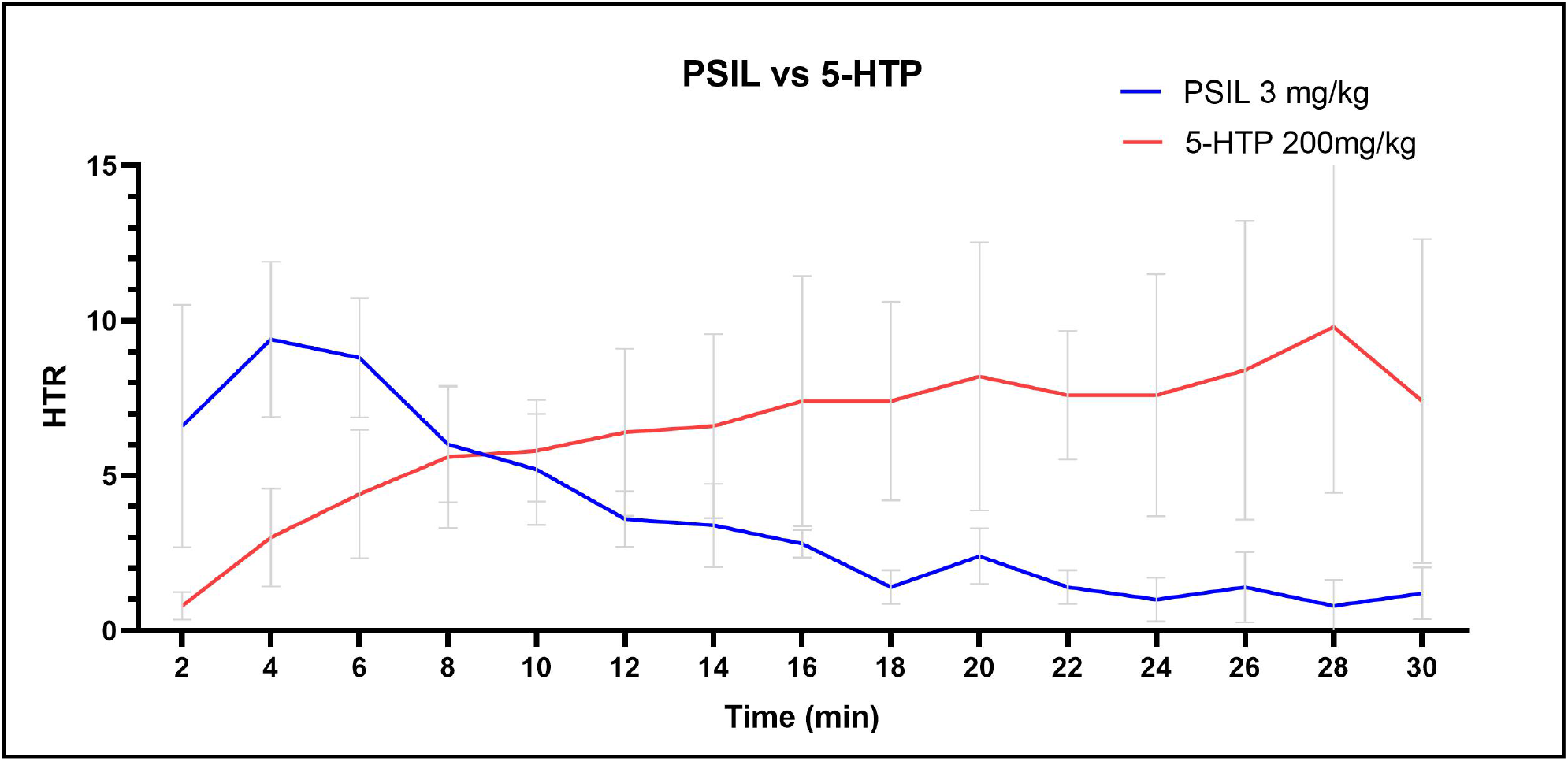
PSIL (3mg/kg) and 5-HTP (200mg/kg) HTR profile comparison based on maximal peak HTR (Twoway ANOVA: Compound F [1, 8] = 6.099, P = 0.0387. Time x Compound F [14, 112] = 13.91, P < 0.0001). Error bars represent SD.

## Discussion

The present study sought to ascertain the effect of 5-HTP and PSIL on HTR over a range of doses and to further our understanding of the receptor mechanisms implicated by pre-treatment administration of receptor modulators. For 5-HTP we defined 200 mg/kg as the highest HTR-inducing dose and demonstrated that both the 5-HT2A antagonist, M100907, and the 5-HT1A agonist, 8-OH-DPAT, significantly blocked the effect. The 5-HT2C antagonist, RS102221, increased 5-HTP-induced HTR up to 4 mg/kg; reduction of HTR was observed with 8 mg/kg. The TAAR1 antagonist EPPTB caused a reduction in 5-HTP-induced HTR with an effective dose of 1 mg/kg. For PSIL we observed a bimodal profile of HTR whereby doses from 1.6 mg/kg elicited a rapid increase in HTR that was followed by a rapid decrease. The higher the administered dose, the more profoundly this effect was demonstrated. Pre-treatment with the 5-HT2A antagonist, M100907, and the 5-HT1A agonist, 8-OH-DPAT, significantly blocked HTR induced by PSIL 4.4 mg/kg. The 5-HT2C antagonist, RS102221, increased PSIL induced HTR up to 4 mg/kg; a significant reduction of HTR was seen with 8 mg/kg. Unlike 5-HTP, there was no effect of the TAAR1 antagonist, EPPTB, on PSIL-induced HTR.

HTR is characterized by a fast, vigorous burst of left to right head-shake movement (González-Maeso et al., 2007; Halberstadt, 2020). Before the development of automated detection methods, manual scoring of HTR was performed, a labor-intensive process that is conducted on one mouse at a time by one scorer. Evolution in the HTR recording field has brought about the use of the magnetometer in conjunction with magnets implanted on the skull of the animals (Halberstadt and Geyer, 2013). In the current study we chose to avoid the surgical procedure for implanting a magnet on the animal’s head and used the novel procedure of attaching magnetic ear tags (de la Fuente Revenga et al., 2020). The magnetic ear tags are well tolerated in mice that are housed in groups. Possible disadvantages can include inflammatory processes that can develop over longer periods of time (during two months after the tagging) and limitations relating to the age and size of the animals. A major advantage is the ability to measure multiple animals as opposed to one when done manually. For the current study, a set-up of 6 individual coiled-containers was used. When using magnetometer recording of HTR, the software used can distinguish between the frequency induced by HTR and the frequency associated with other head movements (de la Fuente Revenga et al., 2019).

An alternative, recently described tool for measuring HTRs in a non-invasive way is the TopScan (Clever Sys Inc, Reston, VA) computer software-based scoring (Glatfelter et al., 2022) system. This system is limited to one mouse per recording box. Another option that was implemented in a recent study is the use of a high-resolution camera recording paired with the software analysis DeepLabCut pro (Mathis Laboratory), which gives video tracking evaluation from a recording but is limited to the number of recordings one can generate as well as false negatives that may arise from the positioning of the mouse during an HTR which might be hidden from the camera and not picked up (Contreras et al., 2021).

5-HTP-induced HTR has previously described by multiple authors (Corne and Pickering, 1967; Darmani and Reeves, 1996; Sun et al., 2003; Devi and Sharma, 2014; Zhuk et al., 2015). However, 5-HTP has not been reported to have psychedelic effects in humans (Turner et al., 2006). Although, overdoses of compounds that increase serotonin release can result in serotonin syndrome which may include hallucinations (Birmes et al., 2003; Evans and Sebastian, 2007), classic psychedelic effects resembling those induced by tryptaminergic and other psychedelic drugs have not been reported. In our study, administration of 5-HTP at 150–250 mg/kg induced significant HTR. The implications of administering equivalent high doses of 5-HTP to humans are unknown. There are two instances of administering up to 3000 mg 5-HTP per os per day but not as a single dose. Such prolonged exposure that can result in tolerance effects (Turner et al., 2006).

Contrasting the induction of HTR between 5-HTP and PSIL, comparison of peak HTR inducing doses of both drugs revealed interesting insights. Firstly, 5-HTP-induced HTR reached a peak at 28 min whereas PSIL induced HTR reached a peak at 4 min. Second, the sharp onset of PSIL that reaches peak HTR during the first 4 min of measurement is associated with a rapid decline, reaching half maximal HTR by 11 min. On the other hand, 5-HTP induced HTR gradually increased over 28 min.

We have categorized the effect on HTR induction by 5-HTP and PSIL of pre-treatment with 5-HT2A and 5-HT2C antagonists and a 5-HT1A agonist. We found complete ablation of HTR induced by 5-HTP and PSIL following pre-treatment with the 5-HT2A antagonist M100907, similar to the effect of M100907 on HTR induced by DOI (Canal et al., 2013; de la Fuente Revenga et al., 2020), 2-CI (Halberstadt and Geyer, 2014), N,N-dipropyltryptamine (DPT) (Fantegrossi et al., 2008), and LSD (Brandt et al., 2016; Rodriguiz et al., 2021; Jaster et al., 2022). The 5-HT1A agonist, 8-OH-DPAT, significantly attenuated HTR induced by 5-HTP and PSIL. Not much literature is available on the effect of 8-OH-DPAT and other 5-HT1A agonists on HTR induced by psychedelic drugs, nevertheless, it has been shown to inhibit DOI-induced head twitch behavior in naive rats (Arnt and Hyttel, 1989; Darmani et al., 1990). In the current study we observed that the 5-HT2C antagonist RS102221 generated an increase in HTR induced by 5-HTP and PSIL up to a dose of 4 mg/kg and decreased HTR with 8 mg/kg. Our observation is comparable to a study that was conducted on the effects of RS102221 on DOI induced HTR (Fantegrossi et al., 2010). Furthermore, DOI-elicited HTR is reduced in 5-HT2C receptor knockout mice (Canal et al., 2010), suggesting the important role that 5-HT2C has in mediating HTR and potentially eliciting psychedelic-like subjective effects in humans. Another recent study found that one of the derivates of DOI (methallylescaline) produced HTR that was blocked by 5-HT2C antagonist (SB-242084), but not by a 5-HT2A antagonist (ketanserin), indicating that the HTR-inducing effect was mediated by 5-HT2C alone (Custodio et al., 2022).

Interestingly, the TAAR1 antagonist EPPTB decreased HTR induced by 5-HTP while not affecting PSIL-induced HTR. There are studies conducted on a broad range of psychoactive drugs regarding their interaction with TAAR1 (Bunzow et al., 2001; Simmler et al., 2016). EPPTB (5 mg/kg) prevented the inhibitory effect of LSD (30–150 μg/kg) on VTA dopamine firing activity (De Gregorio et al., 2016). Although some psychedelics have been found to interact with TAAR1, it seems that there are clear differences between pharmacologically close substances (Wallach, 2009); for example, it has been shown that relative to DMT, 5-MeO-DMT generated a substantially lower TAAR1-induced cAMP response, indicating a lower effectiveness at this receptor (Bunzow et al., 2001). In the same study it has been shown that 5-HTP stimulated a substantial cAMP response at TAAR1 (Bunzow et al., 2001). Current research regarding psilocybin interaction on TAAR1 is interesting, Simmler et al. (2016) have shown that psilocin has Ki values of 1.4 and 17, in rat and mouse TAAR1 respectively. Further studies are needed to clarify the role of TAAR1 receptors in the action of psychedelic drugs.

A significant discussion is under way in the resurgent psychedelics field as to whether short-term subjective effects are needed for the therapeutic potential of psychedelics to be achieved (Olson, 2021; Yaden and Griffiths, 2021). The use of MDMA in the context of psychedelic-assisted psychotherapy has been shown to achieve significant therapeutic effects in PTSD (Mitchell et al., 2021). In the controlled trial of Carhart-Harris (Carhart-Harris et al., 2021) in which psilocybin was shown to be equivalent in efficacy to the SSRI, escitalopram, psychedelic doses of psilocybin were used. Nevertheless, the potential to treat psychiatric disorders without the logistics of trip mediation (use of trained professionals, highly specific space, and duration of effect) has immense economic value. A recent study showed that the attenuation of depression-like features in mice by administration of psilocybin is independent of 5-HT2A receptor activation (demonstrated by co-administrating the 5-HT2A antagonist ketanserin) (Hesselgrave et al., 2021). Our findings suggest the potential use of 5-HT1A agonists to decrease HTR. How such attenuation will impact the therapeutic effect of psychedelics remains to be evaluated. In this context, we have recently demonstrated that co-administration of the 5-HT1A partial agonist, buspirone, which attenuated HTR, did not impede the effect of psilocybin to reduce marble burying (a screening test for anti-obsessional effects) in male ICR mice (Singh et al, Submitted). More research is needed to determine whether attenuation of HTR in mice and of the psychedelic trip in humans by 5-HT receptor modulators, will impede the potential use of psychedelics to treat psychiatric disorders.

## Supporting information

Shahar-HTR-bioRxiv-Supplemental

## Non-standard abbreviations

5-HTP: 5-hydroxytryptophan
5-MeO-DMT: 5-methoxy-N,N-dimethyltryptamine
DMT: N,N-dimethyltryptamine
DOI: 2,5-dimethoxy-4-iodoamphetamine
HTR: head twitch response
i.p.: intraperitoneal
LSD: lysergic acid diethylamide
OCD: obsessive-compulsive disorder
PSIL: chemically synthesized psilocybin
PTSD: posttraumatic disorder
TAAR1: trace amine associated receptor 1

## Acknowledgments

Mario de la Fuente Revenga PhD. of Virginia Commonwealth University provided valuable guidance on the use of the magnetometer apparatus.

Chemical psilocybin was kindly provided by Usona Institute, Madison Wisconsin, USA

Graphical abstract was created with BioRender.com.

## Authorship Contributions

*Participated in research design:* Shahar, Botvinnik, Lifschytz and Lerer.

*Conducted experiments:* Shahar, Botvinnik, Esh-Zuntz, Brownstien, Lifschytz and R. Wolf.

*Contributed new reagents or analytic tools: Esh-Zuntz*

*Performed data analysis:* Shahar, Botvinnik, G. Wolf and Lerer

*Wrote or contributed to the writing of the manuscript:* Shahar, Botvinnik, G. Wolf, Lifschytz and Lerer

## SUPPLEMENTARY MATERIAL

**Supplementary Figure 1.**
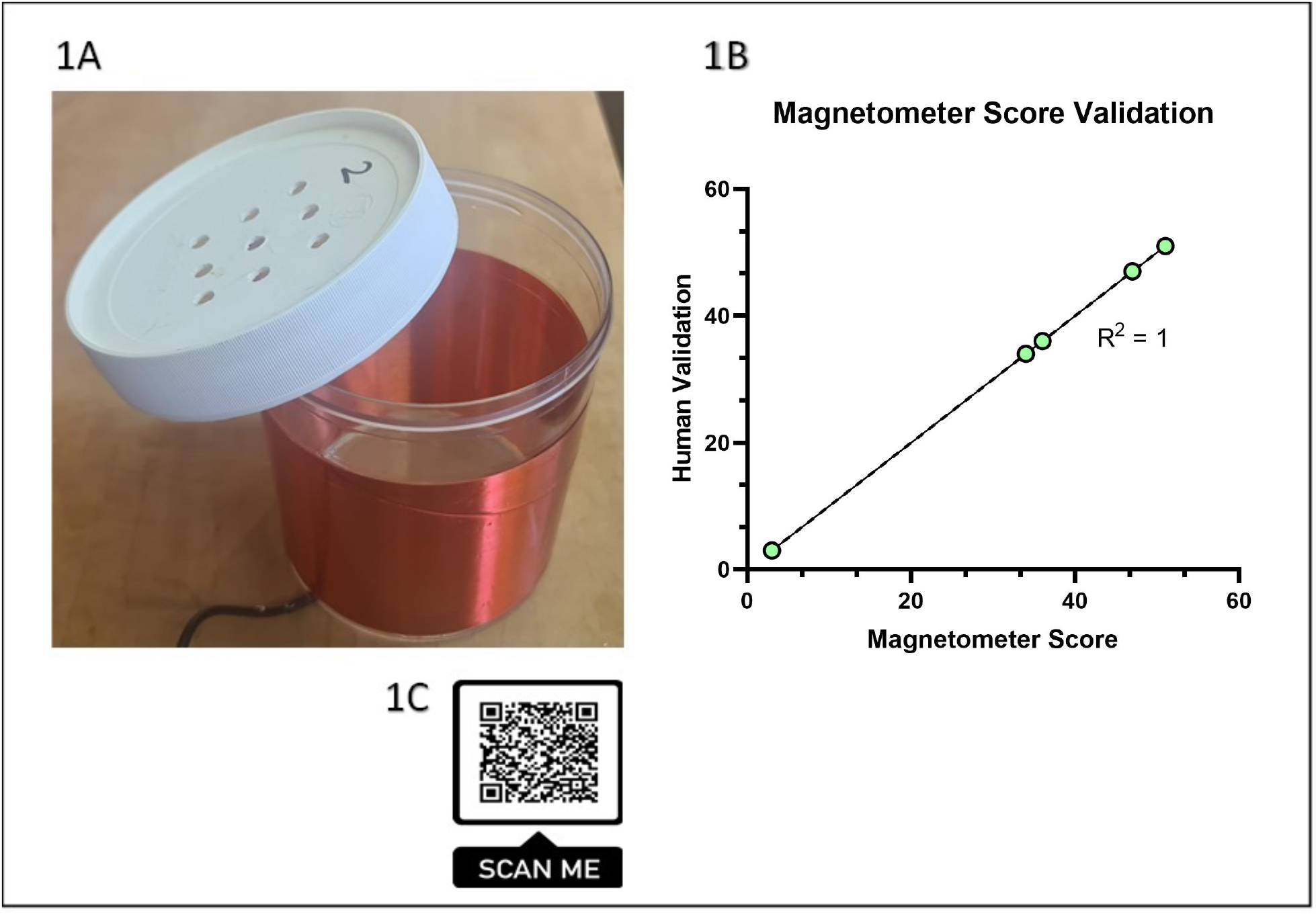
(A) Magnetometer apparatus that was used to measure HTR. One of the six magnetometer coils that make up the system. (B) Validation magnetometer peak scores showed high correlation by human scoring of video of the same mouse while in the magnetometer apparatus (n=5). Simple linear regression: Y = 1*X + 0, R^2^ = 1. F [1,7] = 13892, P < 0.0001. (C) QR code that can be scanned to see a video example of a recorded HTR in the magnetometer.

**Supplementary Movie:**
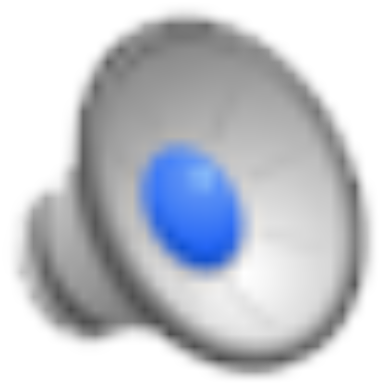
Shows C57Bl/6j mouse manifesting head twitch response after receiving psilocybin 4.4 mg/kg i.p. Each head twitch is immediately preceded by a title. (Filmed by Dr Alexander Botvinnik, Biological Psychiatry Laboratory, Hadassah Medical Center, Hebrew University, Jerusalem, Israel).

## Footnotes

This work was supported in part by Back of the Yards algae sciences (BYAS) and by Parow Entheobiosciences (PEB), Chicago, IL, USA. BL is a consultant to Back of the Yards algae sciences (BYAS) and by Parow Entheobiosciences (PEB) and is listed as a co-inventor on a patent describing concurrent treatment with psychedelics and 5-HT1A agonists that is assigned to BYAS.

## References

Arnt J and Hyttel J (1989) Facilitation of 8-OHDPAT-induced forepaw treading of rats by the 5-HT2 agonist DOI. Eur J Pharmacol 161:45–51.

Birmes P, Coppin D, Schmitt L and Lauque D (2003) Serotonin syndrome: a brief review. CMAJ 168:1439–1442.

Brandt SD, Kavanagh PV, Westphal F, Stratford A, Elliott SP, Hoang K, Wallach J and Halberstadt AL (2016) Return of the lysergamides. Part I: Analytical and behavioural characterization of 1-propionyl-d-lysergic acid diethylamide (1P-LSD). Drug Test Anal 8:891–902.

Brown RT, Nicholas CR, Cozzi NV, Gassman MC, Cooper KM, Muller D, Thomas CD, Hetzel SJ, Henriquez KM, Ribaudo AS and Hutson PR (2017) Pharmacokinetics of Escalating Doses of Oral Psilocybin in Healthy Adults. Clin Pharmacokinet 56:1543–1554.

Bunzow JR, Sonders MS, Arttamangkul S, Harrison LM, Zhang G, Quigley DI, Darland T, Suchland KL, Pasumamula S, Kennedy JL, Olson SB, Magenis RE, Amara SG and Grandy DK (2001) Amphetamine, 3,4-methylenedioxymethamphetamine, lysergic acid diethylamide, and metabolites of the catecholamine neurotransmitters are agonists of a rat trace amine receptor. Mol Pharmacol 60:1181–1188.

Cameron LP, Tombari RJ, Lu J, Pell AJ, Hurley ZQ, Ehinger Y, Vargas MV, McCarroll MN, Taylor JC, Myers-Turnbull D, Liu T, Yaghoobi B, Laskowski LJ, Anderson EI, Zhang G, Viswanathan J, Brown BM, Tjia M, Dunlap LE, Rabow ZT, Fiehn O, Wulff H, McCorvy JD, Lein PJ, Kokel D, Ron D, Peters J, Zuo Y and Olson DE (2021) A non-hallucinogenic psychedelic analogue with therapeutic potential. Nature 589:474–479.

Canal CE, Booth RG and Morgan D (2013) Support for 5-HT2C receptor functional selectivity in vivo utilizing structurally diverse, selective 5-HT2C receptor ligands and the 2,5-dimethoxy-4-iodoamphetamine elicited head-twitch response model. Neuropharmacology 70:112–121.

Canal CE, Olaghere da Silva UB, Gresch PJ, Watt EE, Sanders-Bush E and Airey DC (2010) The serotonin 2C receptor potently modulates the head-twitch response in mice induced by a phenethylamine hallucinogen. Psychopharmacology (Berl) 209:163–174.

Carhart-Harris R, Giribaldi B, Watts R, Baker-Jones M, Murphy-Beiner A, Murphy R, Martell J, Blemings A, Erritzoe D and Nutt DJ (2021) Trial of Psilocybin versus Escitalopram for Depression. N Engl J Med 384:1402–1411.

Carhart-Harris RL and Goodwin GM (2017) The Therapeutic Potential of Psychedelic Drugs: Past, Present, and Future. Neuropsychopharmacology 42:2105–2113.

Castelpietra G, Knudsen AKS, Agardh EE, Armocida B, Beghi M, Iburg KM, Logroscino G, Ma R, Starace F, Steel N, Addolorato G, Andrei CL, Andrei T, Ayuso-Mateos JL, Banach M, Barnighausen TW, Barone-Adesi F, Bhagavathula AS, Carvalho F, Carvalho M, Chandan JS, Chattu VK, Couto RAS, Cruz-Martins N, Dargan PI, Deuba K, da Silva DD, Fagbamigbe AF, Fernandes E, Ferrara P, Fischer F, Gaal PA, Gialluisi A, Haagsma JA, Haro JM, Hasan MT, Hasan SS, Hostiuc S, Iacoviello L, Iavicoli I, Jamshidi E, Jonas JB, Joo T, Jozwiak JJ, Katikireddi SV, Kauppila JH, Khan MAB, Kisa A, Kisa S, Kivimaki M, Koly KN, Koyanagi A, Kumar M, Lallukka T, Langguth B, Ledda C, Lee PH, Lega I, Linehan C, Loureiro JA, Madureira-Carvalho AM, Martinez-Raga J, Mathur MR, McGrath JJ, Mechili EA, Mentis AA, Mestrovic T, Miazgowski B, Mirica A, Mirijello A, Moazen B, Mohammed S, Mulita F, Nagel G, Negoi I, Negoi RI, Nwatah VE, Padron-Monedero A, Panda-Jonas S, Pardhan S, Pasovic M, Patel J, Petcu IR, Pinheiro M, Pollok RCG, Postma MJ, Rawaf DL, Rawaf S, Romero-Rodriguez E, Ronfani L, Sagoe D, Sanmarchi F, Schaub MP, Sharew NT, Shiri R, Shokraneh F, Sigfusdottir ID, Silva JP, Silva R, Socea B, et al. (2022) The burden of mental disorders, substance use disorders and self-harm among young people in Europe, 1990-2019: Findings from the Global Burden of Disease Study 2019. Lancet Reg Health Eur 16:100341.

Cepeda MS, Reps J and Ryan P (2018) Finding factors that predict treatment-resistant depression: Results of a cohort study. Depress Anxiety 35:668–673.

Contreras A, Khumnark M, Hines RM and Hines DJ (2021) Behavioral arrest and a characteristic slow waveform are hallmark responses to selective 5-HT2A receptor activation. Sci Rep 11:1925.

Corne S and Pickering R (1967) A possible correlation between drug-induced hallucinations in man and a behavioural response in mice. Psychopharmacologia 11:65–78.

Corne SJ, Pickering RW and Warner BT (1963) A method for assessing the effects of drugs on the central actions of 5-hydroxytryptamine. Br J Pharmacol Chemother 20:106–120.

Custodio RJP, Ortiz DM, Lee HJ, Sayson LV, Buctot D, Kim M, Lee YS, Kim K-M, Cheong JH and Kim HJ (2022) 5-HT2CR is as important as 5-HT2AR in inducing hallucinogenic effects in serotonergic compounds. Unpublished.

Darmani NA, Martin BR, Pandey U and Glennon RA (1990) Do functional relationships exist between 5-HT1A and 5-HT2 receptors? Journal of Pharmacology Biochemistry Behavior 36:901–906.

Darmani NA and Reeves SL (1996) The mechanism by which the selective 5-HT1A receptor antagonist S-(-)UH 301 produces head-twitches in mice. Pharmacology Biochemistry and Behavior 55:1–10.

De Gregorio D, Posa L, Ochoa-Sanchez R, McLaughlin R, Maione S, Comai S and Gobbi G (2016) The hallucinogen d-lysergic diethylamide (LSD) decreases dopamine firing activity through 5-HT1A, D2 and TAAR1 receptors. Pharmacol Res 113:81–91.

de la Fuente Revenga M, Shah UH, Nassehi N, Jaster AM, Hemanth P, Sierra S, Dukat M and Gonzalez-Maeso J (2021) Psychedelic-like Properties of Quipazine and Its Structural Analogues in Mice. ACS Chem Neurosci 12:831–844.

de la Fuente Revenga M, Shin JM, Vohra HZ, Hideshima KS, Schneck M, Poklis JL and Gonzalez-Maeso J (2019) Fully automated head-twitch detection system for the study of 5-HT2A receptor pharmacology in vivo. Sci Rep 9:14247.

de la Fuente Revenga M, Vohra HZ and Gonzalez-Maeso J (2020) Automated quantification of head-twitch response in mice via ear tag reporter coupled with biphasic detection. J Neurosci Methods 334:108595.

Devi M and Sharma R (2014) Antidepressant activity of aqueous extract of Phaseolus vulgaris (black bean) in rodent models of dep ression. International Journal of Nutrition, Pharmacology, Neurological Diseases 4:118–124.

Evans CE and Sebastian J (2007) Serotonin syndrome. Emergency Medicine Journal 24:e20–e20.

Fantegrossi WE, Harrington AW, Eckler JR, Arshad S, Rabin RA, Winter JC, Coop A, Rice KC and Woods JH (2005) Hallucinogen-like actions of 2,5-dimethoxy-4-(n)-propylthiophenethylamine (2C-T-7) in mice and rats. Psychopharmacology (Berl) 181:496–503.

Fantegrossi WE, Harrington AW, Kiessel CL, Eckler JR, Rabin RA, Winter JC, Coop A, Rice KC and Woods JH (2006) Hallucinogen-like actions of 5-methoxy-N,N-diisopropyltryptamine in mice and rats. Pharmacol Biochem Behav 83:122–129.

Fantegrossi WE, Reissig CJ, Katz EB, Yarosh HL, Rice KC and Winter JC (2008) Hallucinogen-like effects of N,N-dipropyltryptamine (DPT): possible mediation by serotonin 5-HT1A and 5-HT2A receptors in rodents. Pharmacol Biochem Behav 88:358–365.

Fantegrossi WE, Simoneau J, Cohen MS, Zimmerman SM, Henson CM, Rice KC and Woods JH (2010) Interaction of 5-HT2A and 5-HT2C receptors in R(-)-2,5-dimethoxy-4-iodoamphetamine-elicited head twitch behavior in mice. J Pharmacol Exp Ther 335:728–734.

Galvao-Coelho NL, Marx W, Gonzalez M, Sinclair J, de Manincor M, Perkins D and Sarris J (2021) Classic serotonergic psychedelics for mood and depressive symptoms: a meta-analysis of mood disorder patients and healthy participants. Psychopharmacology (Berl) 238:341–354.

Garcia-Romeu A and Richards WA (2018) Current perspectives on psychedelic therapy: use of serotonergic hallucinogens in clinical interventions. Int Rev Psychiatry 30:291–316.

Gershkovich M, Wheaton MG and Simpson HB (2017) Management of Treatment-Resistant Obsessive-Compulsive Disorder. Current Treatment Options in Psychiatry 4:357–370.

Glatfelter GC, Chojnacki MR, McGriff SA, Wang T and Baumann MH (2022) Automated Computer Software Assessment of 5-Hydroxytryptamine 2A Receptor-Mediated Head Twitch Responses from Video Recordings of Mice. ACS Pharmacol Transl Sci 5:321–330.

González-Maeso J, Weisstaub NV, Zhou M, Chan P, Ivic L, Ang R, Lira A, Bradley-Moore M, Ge Y and Zhou Q (2007) Hallucinogens recruit specific cortical 5-HT2A receptor-mediated signaling pathways to affect behavior. Neuron 53:439–452.

González-Maeso J, Yuen T, Ebersole BJ, Wurmbach E, Lira A, Zhou M, Weisstaub N, Hen R, Gingrich JA and Sealfon SC (2003) Transcriptome Fingerprints Distinguish Hallucinogenic and Nonhallucinogenic 5-Hydroxytryptamine 2A Receptor Agonist Effects in Mouse Somatosensory Cortex. The Journal of Neuroscience 23:8836–8843.

Halberstadt AL (2020) Automated detection of the head-twitch response using wavelet scalograms and a deep convolutional neural network. Sci Rep 10:8344.

Halberstadt AL, Chatha M, Klein AK, Wallach J and Brandt SD (2020) Correlation between the potency of hallucinogens in the mouse head-twitch response assay and their behavioral and subjective effects in other species. Neuropharmacology 167:107933.

Halberstadt AL and Geyer MA (2013) Characterization of the head-twitch response induced by hallucinogens in mice: detection of the behavior based on the dynamics of head movement. Psychopharmacology (Berl) 227:727–739.

Halberstadt AL and Geyer MA (2014) Effects of the hallucinogen 2,5-dimethoxy-4-iodophenethylamine (2C-I) and superpotent N-benzyl derivatives on the head twitch response. Neuropharmacology 77:200–207.

Hesselgrave N, Troppoli TA, Wulff AB, Cole AB and Thompson SM (2021) Harnessing psilocybin: antidepressant-like behavioral and synaptic actions of psilocybin are independent of 5-HT2R activation in mice. Proc Natl Acad Sci U S A 118.

Hofmann A, Heim R, Brack A, Kobel H, Frey A, Ott H, Petrzilka T and Troxler F (1959) Psilocybin und Psilocin, zwei psychotrope Wirkstoffe aus mexikanischen Rauschpilzen. Helvetica Chimica Acta 42:1557–1572.

Janhavi P, Divyashree S, Sanjailal KP and Muthukumar SP (2022) DoseCal: a virtual calculator for dosage conversion between human and different animal species. Arch Physiol Biochem 128:426–430.

Jaster AM, Elder H, Marsh SA, de la Fuente Revenga M, Negus SS and Gonzalez-Maeso J (2022) Effects of the 5-HT2A receptor antagonist volinanserin on head-twitch response and intracranial self-stimulation depression induced by different structural classes of psychedelics in rodents. Psychopharmacology (Berl) 239:1665–1677.

Keiser MJ, Setola V, Irwin JJ, Laggner C, Abbas AI, Hufeisen SJ, Jensen NH, Kuijer MB, Matos RC, Tran TB, Whaley R, Glennon RA, Hert J, Thomas KL, Edwards DD, Shoichet BK and Roth BL (2009) Predicting new molecular targets for known drugs. Nature 462:175–181.

Knapp M and Wong G (2020) Economics and mental health: the current scenario. World Psychiatry 19:3–14.

Krediet E, Bostoen T, Breeksema J, van Schagen A, Passie T and Vermetten E (2020) Reviewing the Potential of Psychedelics for the Treatment of PTSD. Int J Neuropsychopharmacol 23:385–400.

Kyzar EJ, Nichols CD, Gainetdinov RR, Nichols DE and Kalueff AV (2017) Psychedelic Drugs in Biomedicine. Trends Pharmacol Sci 38:992–1005.

Lu J, Tjia M, Mullen B, Cao B, Lukasiewicz K, Shah-Morales S, Weiser S, Cameron LP, Olson DE, Chen L and Zuo Y (2021) An analog of psychedelics restores functional neural circuits disrupted by unpredictable stress. Mol Psychiatry 26:6237–6252.

Mitchell JM, Bogenschutz M, Lilienstein A, Harrison C, Kleiman S, Parker-Guilbert K, Ot’alora GM, Garas W, Paleos C, Gorman I, Nicholas C, Mithoefer M, Carlin S, Poulter B, Mithoefer A, Quevedo S, Wells G, Klaire SS, van der Kolk B, Tzarfaty K, Amiaz R, Worthy R, Shannon S, Woolley JD, Marta C, Gelfand Y, Hapke E, Amar S, Wallach Y, Brown R, Hamilton S, Wang JB, Coker A, Matthews R, de Boer A, Yazar-Klosinski B, Emerson A and Doblin R (2021) MDMA-assisted therapy for severe PTSD: a randomized, double-blind, placebo-controlled phase 3 study. Nat Med 27:1025–1033.

Nichols DE (2016) Psychedelics. 68:264–355.

Nutt D (2022) Psychedelic drugs—a new era in psychiatry? Dialogues in clinical neuroscience.

Olbrich S, Preller KH and Vollenweider FX (2021) LSD and ketanserin and their impact on the human autonomic nervous system. Psychophysiology 58:e13822.

Olson DE (2021) Are the Subjective Effects of Psychedelics Necessary for Their Enduring Therapeutic Effects? A conversation with David E. Olson and David B. Yaden. The editors wish to express their gratitude to ALIUS members for their valuable help in proofreading the interviews, and to all the contributors for accepting to participate to this issue.

Passie T, Seifert J, Schneider U and Emrich HM (2002) The pharmacology of psilocybin. Addict Biol 7:357–364.

Pokorny T, Preller KH, Kraehenmann R and Vollenweider FX (2016) Modulatory effect of the 5-HT1A agonist buspirone and the mixed non-hallucinogenic 5-HT1A/2A agonist ergotamine on psilocybin-induced psychedelic experience. Eur Neuropsychopharmacol 26:756–766.

Rodriguiz RM, Nadkarni V, Means CR, Chiu Y-T, Roth BL and Wetsel WC (2021) LSD’s effects are differentially modulated in arrestin knockout mice.2021.2002.2004.429772.

Siegel AN, Meshkat S, Benitah K, Lipsitz O, Gill H, Lui LMW, Teopiz KM, McIntyre RS and Rosenblat JD (2021) Registered clinical studies investigating psychedelic drugs for psychiatric disorders. J Psychiatr Res 139:71–81.

Simmler LD, Buchy D, Chaboz S, Hoener MC and Liechti ME (2016) In vitro characterization of psychoactive substances at rat, mouse, and human trace amine-associated receptor 1. Journal of Pharmacology Experimental Therapeutics 357:134–144.

Slocum ST, DiBerto JF and Roth BL (2022) Molecular insights into psychedelic drug action. J Neurochem 162:24–38.

Sun H-L, Zheng J-W, Wang K, Liu R-K and Liang J-H (2003) Tramadol reduces the 5-HTP-induced head-twitch response in mice via the activation of μ and κ opioid receptors. Life Sciences 72:1221–1230.

Turner EH, Loftis JM and Blackwell AD (2006) Serotonin a la carte: supplementation with the serotonin precursor 5-hydroxytryptophan. Pharmacol Ther 109:325–338.

van Amsterdam J and van den Brink W (2022) The therapeutic potential of psilocybin: a systematic review. Expert Opin Drug Saf:1–8.

Vollenweider FX, Vollenweider-Scherpenhuyzen MF, Babler A, Vogel H and Hell D (1998) Psilocybin induces schizophrenia-like psychosis in humans via a serotonin-2 agonist action. Neuroreport 9:3897–3902.

Wallach J (2009) Endogenous hallucinogens as ligands of the trace amine receptors: a possible role in sensory perception. Medical hypotheses 72:91–94.

Winkelman M (2019) Introduction: Evidence for entheogen use in prehistory and world religions. Journal of Psychedelic Studies 3:43–62.

Yaden DB and Griffiths RR (2021) The Subjective Effects of Psychedelics Are Necessary for Their Enduring Therapeutic Effects. ACS Pharmacol Transl Sci 4:568–572.

Zhuk O, Jasicka-Misiak I, Poliwoda A, Kazakova A, Godovan VV, Halama M and Wieczorek PP (2015) Research on acute toxicity and the behavioral effects of methanolic extract from psilocybin mushrooms and psilocin in mice. Toxins (Basel) 7:1018–1029.

