## Supplementary material for "Role of 5-HT2A, 5-HT2C, 5-HT1A and TAAR1 receptors in the head twitch response induced by 5-hydroxytryptophan and psilocybin: Translational implications": Shahar-HTR-bioRxiv-Supplemental

**Shahar et al**

**SUPPLEMENTARY MATERIAL**

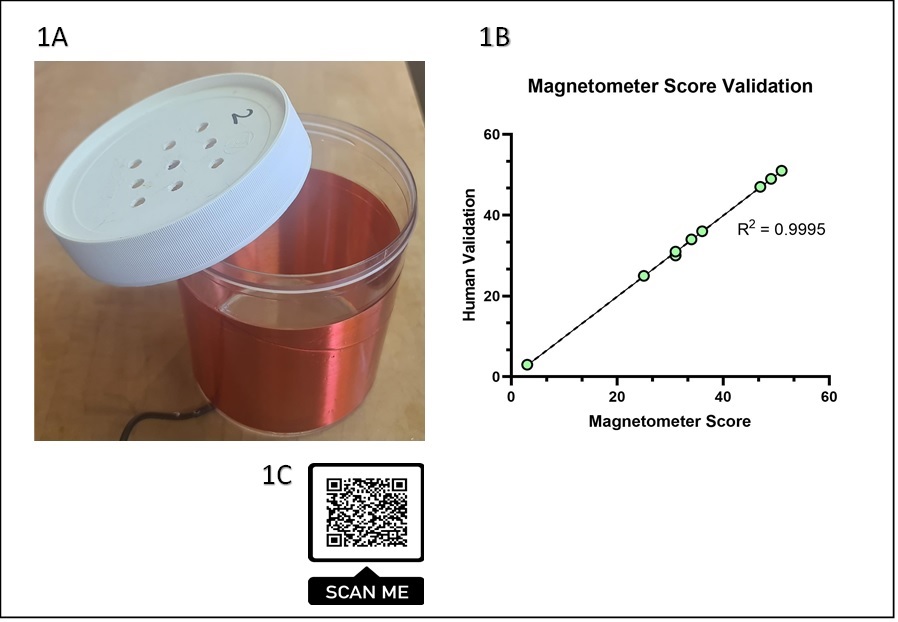

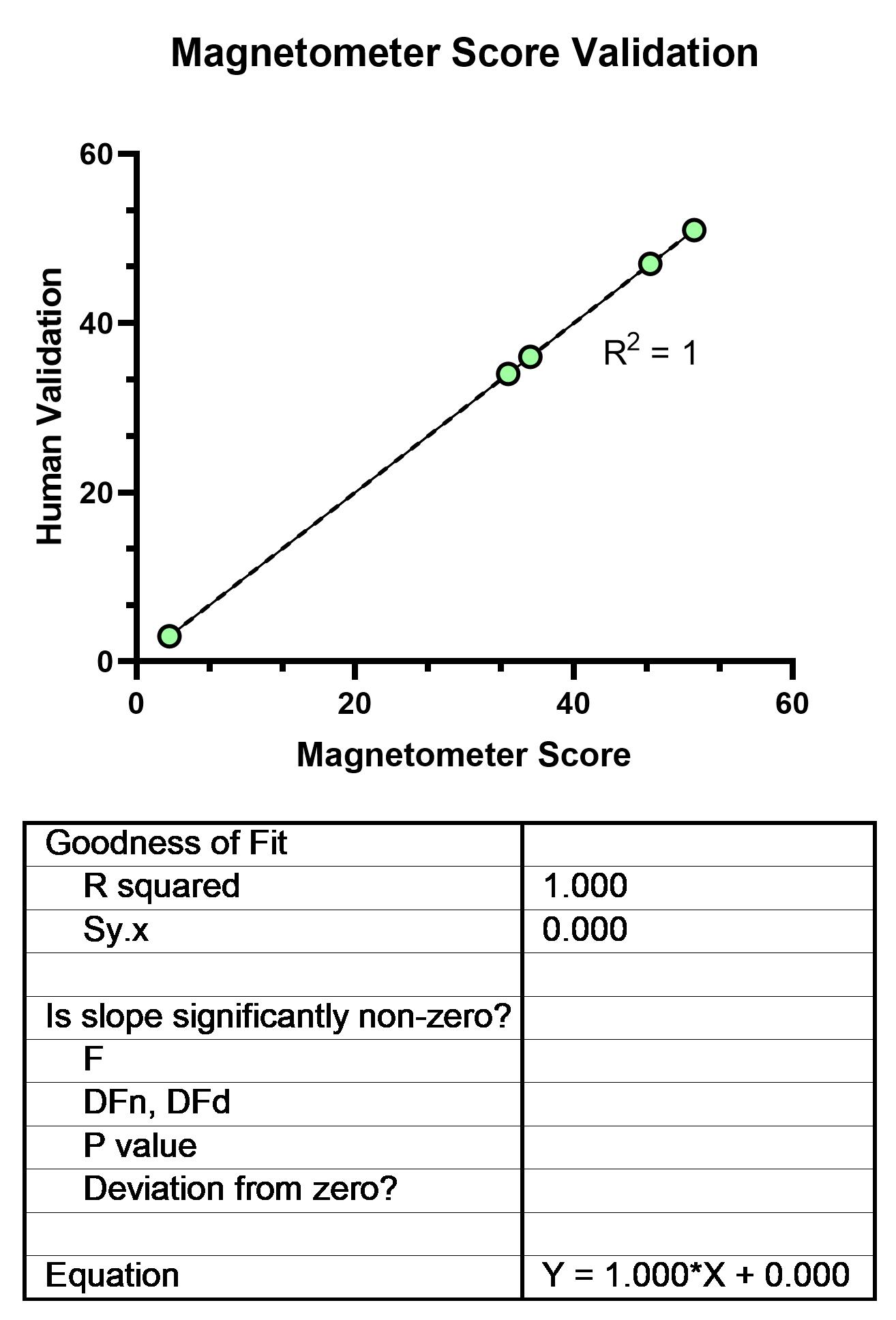

**Supplementary Figure 1**. (A) Magnetometer apparatus that was used to measure HTR. One of the six magnetometer coils that make up the system. (B) Validation magnetometer peak scores showed high correlation by human scoring of video of the same mouse while in the magnetometer apparatus (n=5). Simple linear regression: Y = 1*X + 0, R^2^ = 1. F [1,7] = 13892, P < 0.0001. (C) QR code that can be scanned to see a video example of a recorded HTR in the magnetometer.

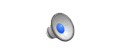

**Supplementary Movie:** Shows C57Bl/6j mouse manifesting head twitch response after receiving psilocybin 4.4 mg/kg i.p. Each head twitch is immediately preceded by a title. (Filmed by Dr Alexander Botvinnik, Biological Psychiatry Laboratory, Hadassah Medical Center, Hebrew University, Jerusalem, Israel).
